# Distinct D-box Motifs in SPD-2 Mediate APC/C^FZR-1^-Dependent Degradation and Centrosomal Localization in *Caenorhabditis elegans* Embryos

**DOI:** 10.1101/2025.10.14.682142

**Authors:** Rachel N. Yim, Joseph R. DiPanni, Paollette A. Rivera, Mi Hye Song

## Abstract

Centrosome duplication must be tightly regulated to maintain genomic stability. In *Caenorhabditis elegans*, the APC/C and co-activator FZR-1 function as negative regulators of centrosome duplication by targeting specific substrates for proteolytic degradation. While *C. elegans* SAS-5 and ZYG-1 have been identified as substrates of APC/C^FZR-1^, the mechanism by which APC/C^FZR-1^-dependent degradation influences centrosome assembly remains unclear. Here, we identified SPD-2, the conserved homolog of human CEP192, as a substrate of APC/C^FZR-1^. We show that loss of APC/C^FZR-1^ increases both cellular and centrosomal SPD-2 levels, and that SPD-2 physically associates with FZR-1 *in vivo*. Functional analyses of canonical D-box motifs reveal that D-box1, D-box2, and D-box3 each contribute to SPD-2 degradation, each with different functional consequences. Mutation of D-box3 alone partially rescued *zyg-1* mutant phenotypes by restoring centrosome duplication and embryonic viability through increased centrosomal SPD-2 and ZYG-1. In contrast, mutating D-box1 or D-box2 elevated cellular SPD-2 but did not rescue *zyg-1*, with the D-box1 mutation further reducing centrosomal SPD-2 and exacerbating duplication defects and lethality in *zyg-1* mutants. Our results reveal a conserved mechanism for APC/C^FZR-1^-dependent degradation of SPD-2 and show that its degron motifs have dual functions in degradation and centrosomal localization, ensuring robust control of centrosome assembly during *C. elegans* embryogenesis.

## Introduction

Centrosomes are the primary microtubule-organizing centers (MTOCs) in animal cells and play crucial roles in spindle assembly, chromosome segregation, and cell fate determination (Nigg and Holland 2018). Their functional capacity depends on the dynamic recruitment of pericentriolar material (PCM) around a centriole core, ensuring that centrosomes are competent for duplication and microtubule nucleation during each cell cycle. Errors in centrosome regulation can lead to defective mitotic spindles, genomic instability, and developmental disorders, including microcephaly and cancer (Fernandes-Mariano et al. 2025). Understanding how centrosome assembly and activity are coordinated in both time and space is therefore crucial in cell and developmental biology.

A key step in centrosome biogenesis and maturation in *Caenorhabditis elegans* is the recruitment of centrosome components to the site of new centriole formation, a process initiated by the conserved scaffold protein SPD-2 (Kemp et al. 2004; Pelletier et al. 2004) and its upstream factor SAS-7 (Sugioka et al. 2017). SPD-2 localizes to both centrioles and PCM, where it acts upstream of the kinase ZYG-1 (Pelletier et al. 2006; Shimanovskaya et al. 2014) and promotes the recruitment of SPD-5 and γ-tubulin to link PCM assembly with centriole duplication (Kemp et al. 2004; Pelletier et al. 2004). Loss of SPD-2 function significantly impairs centrosome biogenesis, maturation, and microtubule dynamics, underscoring its vital role in centrosome assembly and function (Kemp et al. 2004; Pelletier et al. 2004). Recent structural analyses have further uncovered the modular and intrinsically disordered structure of SPD-2, highlighting its capacity to mediate protein–protein interactions that link centrosome duplication with PCM expansion (Murph et al. 2022). Although SPD-2 is known to be crucial for centrosome regulation, its abundance and activity during centrosome assembly in *C. elegans* embryos remain unclear.

Proteolytic regulation by the anaphase-promoting complex/cyclosome (APC/C), an E3 ubiquitin ligase, has emerged as a key mechanism for controlling centrosome activity and maintaining centrosome copy number (Hames et al. 2001; Strnad et al. 2007; Arquint and Nigg 2014; Meghini et al. 2016; Medley et al. 2017; Medley et al. 2021; Tischer et al. 2022; Meghini et al. 2023). In *Drosophila*, the APC/C, in complex with its co-activator Fizzy-related (Fzr), targets Spd2 for degradation (Meghini et al. 2016). This degradation maintains centrosome asymmetry by limiting Spd2 accumulation at the daughter centrosome, ensuring proper PCM recruitment and maintaining spindle orientation. Without APC/C^Fzr^-dependent proteolysis, excess Spd2 disrupts centrosome behavior, impairs microtubule nucleation, and randomizes the division axis (Meghini et al. 2023). These findings demonstrate that proteasome-mediated turnover of conserved centrosome proteins is a crucial mechanism for maintaining centrosome function. Given the conserved role of SPD-2 in centrosome assembly, a similar APC/C-dependent regulation may occur in *C. elegans*. Indeed, SPD-2 functions upstream of ZYG-1 in the centrosome assembly pathway, indicating that precise regulation of SPD-2 levels is critical for promoting centrosome duplication and coordinating centrosome maturation (Kemp et al. 2004; Pelletier et al. 2004). Dysregulation of SPD-2 levels could therefore have serious consequences for centrosome assembly and function during the cell cycle.

In *C. elegans*, it has been shown that APC/C^FZR-1^ regulates centrosome duplication by promoting the degradation of the centrosome protein SAS-5 through its KEN-box motif (Medley et al. 2017) and by modulating centrosomal ZYG-1 levels via D-box motifs (Medley et al. 2021). Loss of FZR-1 or core APC/C subunits elevates SAS-5 and ZYG-1 levels at centrosomes, thereby restoring bipolar spindle formation in hypomorphic *zyg-1* mutants, revealing that APC/C^FZR-1^ functions at multiple points in the centrosome assembly pathway (Medley et al. 2017; Medley et al. 2021). Together, these genetic analyses indicate that APC/C^FZR-1^ targets multiple substrates, including ZYG-1 and SAS-5, as well as additional factors acting upstream of ZYG-1. Here, we investigate the hypothesis that APC/C^FZR-1^-mediated degradation of SPD-2 provides a conserved mechanism to regulate centrosome assembly in *C. elegans* embryos.

## Materials and Methods

### *C. elegans* Culture and Genetic Analysis

The *C. elegans* strains used in this study are listed in Supplementary **Table 1**. All strains were derived from the wild-type Bristol N2 strain (Brenner 1974; Church et al. 1995) and maintained on MYOB plates seeded with Escherichia coli OP50 at 16°C or 20°C. Some strains were provided by the CGC, which is funded by NIH Office of Research Infrastructure Programs (P40 OD010440). For genetic analysis, individual L4 hermaphrodites were transferred to new plates and allowed to produce progeny for 0-24, 24-48, and 48-72 hours at the indicated temperatures. Progeny were allowed to develop for 18-24 hours before counting the number of larvae and dead eggs.

### Immunofluorescence and Cytological analysis

Immunofluorescence and confocal microscopy were performed as described (Stubenvoll et al. 2016). The following primary and secondary antibodies were used at 1:3000 dilutions: DM1a (Millipore Sigma, #T9026), α-ZYG-1 (Stubenvoll et al. 2016), α-TBG-1 (Stubenvoll et al. 2016), α-HA (Millipore Sigma, #11867423001, #11583816001; Thermo Fisher, #26183-A647), Alexa Fluor 488 and 568 secondary antibodies (Thermo Fisher, #A11001, A11004, A11006, A11034, A11036). Confocal microscopy was performed using a Nikon Eclipse Ti-U microscope equipped with a Plan Apo 60×1.4 NA lens, a Spinning Disk Confocal (CSU X1), and a Photometrics Evolve 512 camera. MetaMorph software (Molecular Devices, Sunnyvale, CA, USA) was employed for image acquisition and fluorescence intensity quantification, and Adobe Photoshop/Illustrator 2025 were used for image processing. To quantify fluorescent signals at centrosomes, the average intensity within an 8- or 9-pixel (1 pixel = 0.151 µm) diameter region was recorded from the highest-intensity focal plane within an area centered on each centrosome. The average intensity within a 25-pixel diameter region outside the embryo was used for background subtraction.

### CRISPR/Cas9 Genome Editing

For genome editing, we used the co-CRISPR method described previously (Arribere et al. 2014; Paix et al. 2015). crRNA was designed using the CRISPOR web server (crispor.tefor.net; Concordet and Haeussler, 2018). Animals were microinjected with a mixture of commercially available SpCas9 (IDT, Coralville, IA) and custom-designed oligonucleotides, including crRNAs (Supplementary **Table 2**) at 0.4–0.8 mg/ml, tracrRNA at 12 mg/ml, and single-stranded DNA oligonucleotides (Supplementary **Table 3**) at 25–100 ng/ml. The amount of crRNA was tripled for low-efficiency crRNAs (SPD-2:D-box2). After injection, we screened for *dpy-10(cn64) II/+* rollers in F1 progeny and genotyped F2. The genome editing was confirmed by Sanger sequencing (GeneWiz, South Plainfield, NJ).

### Immunoprecipitation (IP) and Immunoblot

IP experiments were performed as described previously (Medley et al. 2021). Embryos were collected from young gravid worms using hypochlorite treatment (1:2:1 ratio of M9 buffer, 5.25% sodium hypochlorite, and 5 M NaOH), frozen in liquid nitrogen, and stored at -80°C until use. Embryos were suspended in lysis buffer [50 mM HEPES, pH 7.4, 2 mM EDTA, pH 8.0, 1 mM MgCl_2_, 150 mM KCl, 0.5 mM DTT, 0.5% NP-40 (v/v) and 10% glycerol (v/v)] with complete protease inhibitor cocktail (Millipore Sigma) and MG132 (Tocris, Avonmouth, Bristol, UK), milled for 8 minutes (repeat x2) at 30 Hz using a Retsch MM 400 mixer-mill (Verder Scientific, Newtown, PA), then sonicated for 5 minutes in an ultrasonic bath (Thermo Fisher). Lysates were spun at 45,000 rpm for 45 minutes using a Sorvall RC M120EX ultracentrifuge (Thermo Fisher), and the supernatant was recovered into clean tubes. An equal amount of total protein lysates was used for each IP. Lysates and α-HA tag magnetic beads (MBL, #M181-11; Millipore Sigma, #SAE0197) were incubated with rotation for 1 hour at 4°C and washed (5x 5 minutes) with PBST (PBS + 0.1% Tween-20). IP with beads and input samples were resuspended in 2X Laemmli Sample Buffer (Sigma) and boiled for 5 minutes before fractionation on a 4–12% NuPAGE Bis-Tris gel (Thermo Fisher Scientific). Proteins on a gel were transferred to a nitrocellulose membrane and analyzed using the antibodies at 1:3000-10,000 dilutions: α-SPD-2 (Song et al., 2008), α-TBG-1 (Stubenvoll et al., 2016), DM1a (Sigma, #T9026), α-Ollas (Thermo Fisher, #MA5-16125), α-HA (Millipore Sigma, #11867423001, #11583816001), IRDye secondary antibodies (LI-COR Biosciences). Blots were imaged with the Odyssey M scanner (LI-COR Biosciences) and analyzed using Image Studio software (LI-COR Biosciences).

### Statistical analysis

Statistics were produced using R statistical software and presented as average ± standard deviation (SD). Dot plots were generated using the R ‘beeswarm’ package. In the dotplots, the box ranges from the first through the third quartile of the data. The thick bar indicates the median. A solid grey line extends 1.5 times the interquartile range, or to the minimum and maximum data points. All P-values were calculated using two-tailed t-tests: ^ns^ *p*>0.05, * *p*<0.05, ** *p*<0.01, *** *p*<0.001.

## Results and Discussion

### Loss of FZR-1 stabilizes cellular SPD-2

In *C. elegans* embryos, APC/C^FZR-1^ negatively regulates centrosome duplication by promoting proteasomal degradation of SAS-5 (Medley et al. 2017) and by modulating centrosomal ZYG-1 levels (Medley et al. 2021). Previous genetic analyses indicate that APC/C^FZR-1^ targets multiple substrates, including ZYG-1 and SAS-5, as well as other factors acting upstream of ZYG-1 (Medley et al. 2021). Here, we investigate SPD-2, which functions upstream of ZYG-1 in the centrosome assembly pathway, as a potential substrate of APC/C^FZR-1^ in *C. elegans* embryos. If APC/C^FZR-1^ directly targets SPD-2 for proteasomal degradation, then inhibiting APC/C^FZR-1^ should block SPD-2 degradation, thereby hyper-stabilizing SPD-2 and elevating its overall cellular levels. We first tested whether loss of APC/C^FZR-1^ affected SPD-2 abundance by quantitative Western blot analysis using anti-SPD-2 (Song et al. 2008) and protein lysates extracted from *fzr-1(bs31)* mutant (Medley et al. 2017) and wild-type (WT) embryos (**Fig. 1A**). Our quantification data showed that SPD-2 levels in *fzr-1(bs31)* mutant embryos are significantly higher (1.89±0.71-fold, *p*<0.001, n=27) compared to WT controls (1.00±0.0-fold, n=24), while *zyg-1* mutant embryos show no significant change (0.94±0.16-fold, n=10). In contrast, TBG-1 levels appear unaffected in both mutants. We also observed a similar trend in *mat-3(or180)* for the core APC8/CDC23 subunit (Golden et al. 2000) and *emb-1(hc57)* mutants for the APC16 (Shakes et al. 2011) in the *C. elegans* APC/C complex (Supplementary **Fig. 1A**). These findings indicate that loss of APC/C^FZR-1^ results in elevated cellular SAS-5 levels, most likely due to increased protein stability caused by impaired proteasomal degradation.

**Figure 1.**
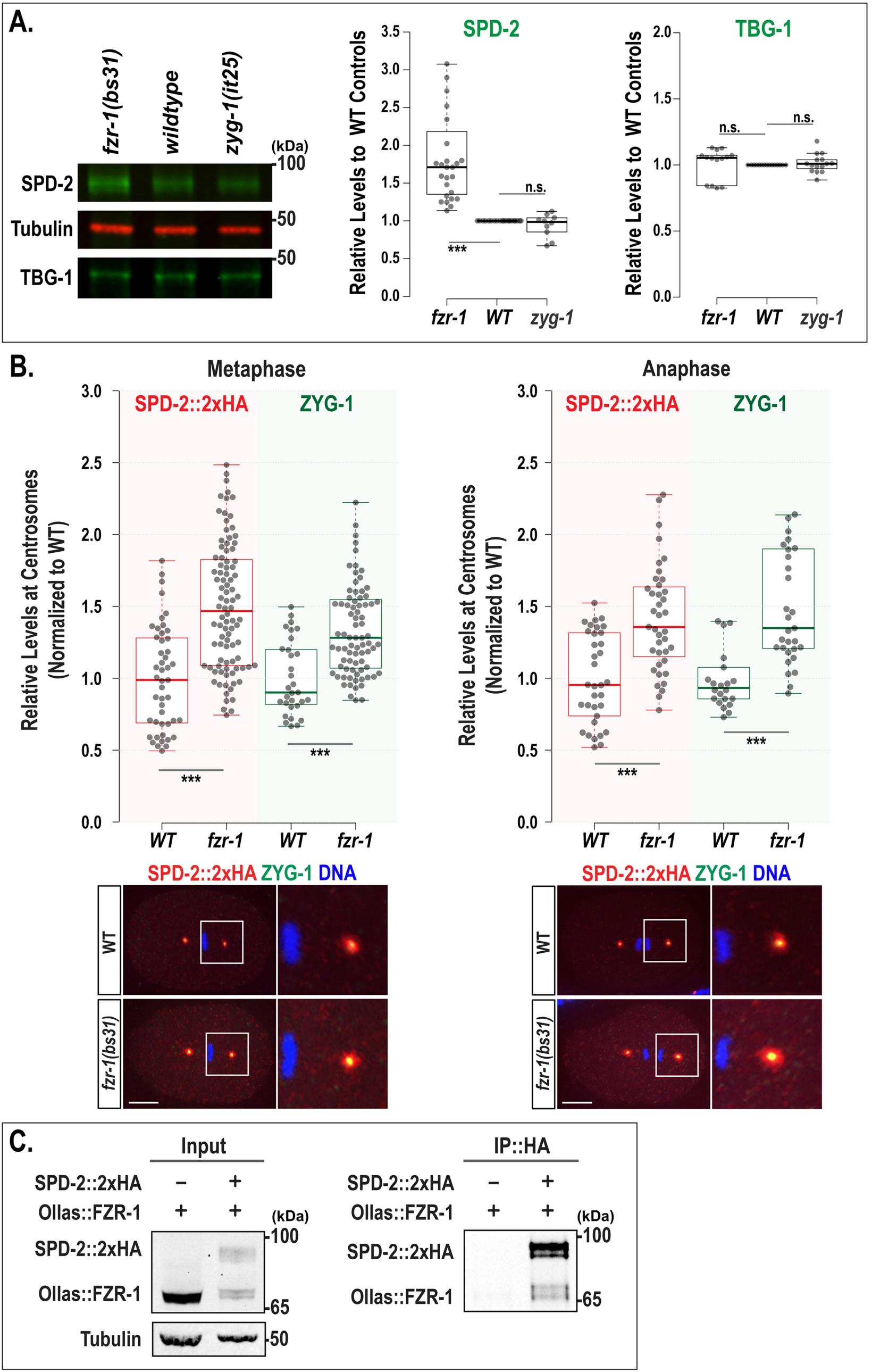
Loss of FZR-1 stabilizes SPD-2. (**A**) Representative western blot with anti-SPD-2 and anti-TBG-1 using embryonic lysates extracted from *fzr-1(bs31)* mutants, wild-type(*N2*), and *zyg-1(it25)* mutants (Left). Tubulin was used as a loading control. Quantitative Western blot analyses show significantly increased levels of SPD-2 (Middle) in *fzr-1(bs31)* mutants (1.89±0.71-fold, *p*<0.001, n=27), but no change in *zyg-1(it25)* mutants (0.94±0.16-fold, *p*=0.49, n=10), compared to wild-type (WT) controls (1.00±0.0-fold, n=24). In contrast, there were no significant differences in TBG-1 levels (Right) among *fzr-1(bs31)* mutants (1.01±0.12-fold, *p=*0.80, n=14), *zyg-1(it25)* mutants (1.01±0.07-fold, *p=*0.53, n=15), and wild-type controls (1.00±0.0-fold, n=15). (**B**) (Top) Quantification of total centrosomal SPD-2::2xHA and ZYG-1 levels during the first mitotic cycle. Each dot represents a centrosome. (Bottom) Embryos stained for SPD-2::2xHA and ZYG-1 at the first mitotic metaphase and anaphase, with magnified images of posterior centrosomes. Bars, 10 μm. In the plots, the box ranges from the first through the third quartile of the data. The thick bar indicates the median. (**A-B**) The solid grey line extends 1.5 times the interquartile range or to the minimum and maximum data points. ^ns^*p*>0.05, \*\*\**p*<0.001 (two-tailed t-tests). (**C**) IP using anti-HA suggests that SPD-2 physically interacts with FZR-1. Protein lysates extracted from embryos expressing only Ollas::FZR-1 were used as a negative control; ∼1% of the total lysates were loaded in the input lanes. Tubulin was used as a loading control in the input lanes.

Then, increased levels of cellular SPD-2 in the embryo can influence the amount of SPD-2 recruited to centrosomes. To assess SPD-2 levels at centrosomes, we stained embryos with anti-HA and measured the fluorescence intensity of the SPD-2::2xHA signal at centrosomes (Medley et al. 2023). As SPD-2 localizes to both centrioles and PCM, we measured total centrosomal (centriolar and PCM; **Fig. 1B**) and only centriolar SPD-2 signals (Supplementary **Fig. 1C**). Quantitative immunofluorescence (IF) using SPD-2 tagged with 2xHA (SPD-2::2xHA) reveal that during the first mitosis, *fzr-1(bs31)* mutant embryos exhibit increased levels of total centrosomal SPD-2::2xHA (1.50±0.44-fold, n=92 at metaphase; 1.42±0.37-fold, n=39 at anaphase) compared to WT controls (1.00±0.36-fold, n=47 at metaphase; 1.00±0.31-fold, n=33 at anaphase). A comparable increase was seen in *mat-3(or180)* for the core APC8/CDC23 subunit (Golden et al. 2000) (Supplementary **Fig. 1B**). Additionally, a similar pattern was observed for the centriolar SPD-2 levels (Supplementary **Fig. 1C**). The *fzr-1(bs31)* mutant embryo exhibits increased levels of centriolar SPD-2::2xHA (1.50±0.37-fold, n=98 at metaphase; 1.36±0.44-fold, n=39 at anaphase) compared to WT controls (1.00±0.19-fold, n=34 at metaphase; 1.00±0.30-fold, n=36 at anaphase). Since our prior study showed that *fzr-1(bs31)* mutant embryos display elevated centrosomal ZYG-1 levels (Medley et al. 2021), we also measured ZYG-1 levels by co-staining embryos with anti-HA and anti-ZYG-1 (Stubenvolle et al. 2016; **Fig. 1B**). Consistent with prior observations, *fzr-1(bs31)* mutant embryos exhibit increased levels of centrosomal ZYG-1 (1.35±0.35-fold, n=80 at metaphase; 1.55±0.49-fold, n=31 at anaphase) compared to WT controls (1.00±0.26-fold, n=31 at metaphase; 1.00±0.23-fold, n=32 at anaphase). Our data show that loss of APC/C^FZR-1^ leads to a significant increase in both cellular and centrosomal levels of SPD-2, supporting a model in which APC/C^FZR-1^-dependent proteolysis regulates SPD-2 abundance during *C. elegans* embryogenesis.

Furthermore, our immunoprecipitation (IP) assays demonstrate a physical interaction between SPD-2 and FZR-1 *in vivo* (**Fig. 1C**). We detected SPD-2::2xHA and Ollas::FZR-1 in the pull-down using embryonic lysates that express SPD-2::2xHA (Medley et al. 2023) and Ollas::FZR-1 (Medley et al. 2021). These results collectively support our hypothesis that APC/C^FZR-1^ directly targets SPD-2 for proteasomal degradation. These findings support our model in which the APC/C and its co-activator FZR-1 (APC/C^FZR-1^) control SPD-2 levels in *C. elegans* embryos.

### SPD-2 contains five putative D-box motifs

The E3 ubiquitin ligase APC/C^FZR-1^ targets specific substrates through the coactivator FZR-1, which directly recognizes conserved motifs such as the destruction (D)-box and KEN-box within the substrates (Glotzer et al. 1991; Pfleger and Kirschner 2000). Our *in silico* analysis identified five potential D-box motifs (RxxL), but no KEN-box, within the *C. elegans* SPD-2 (**Fig. 2A**). In SPD-2, D-box1 is located within the coiled-coil (CC3: aa291-322) domain, D-box2 within the Aspm-SPD-2-Hydin (ASH) domain, D-box3 between the Ig-like and PDZ domains, and both D-box4 and D-box5 within the PDZ domain (Murph et al. 2022). The sequence alignments indicate that D-box4 is the only fully conserved motif across closely related nematodes, while the other motifs retain partial but detectable sequence conservation (**Fig. 2B**).

**Figure 2.**
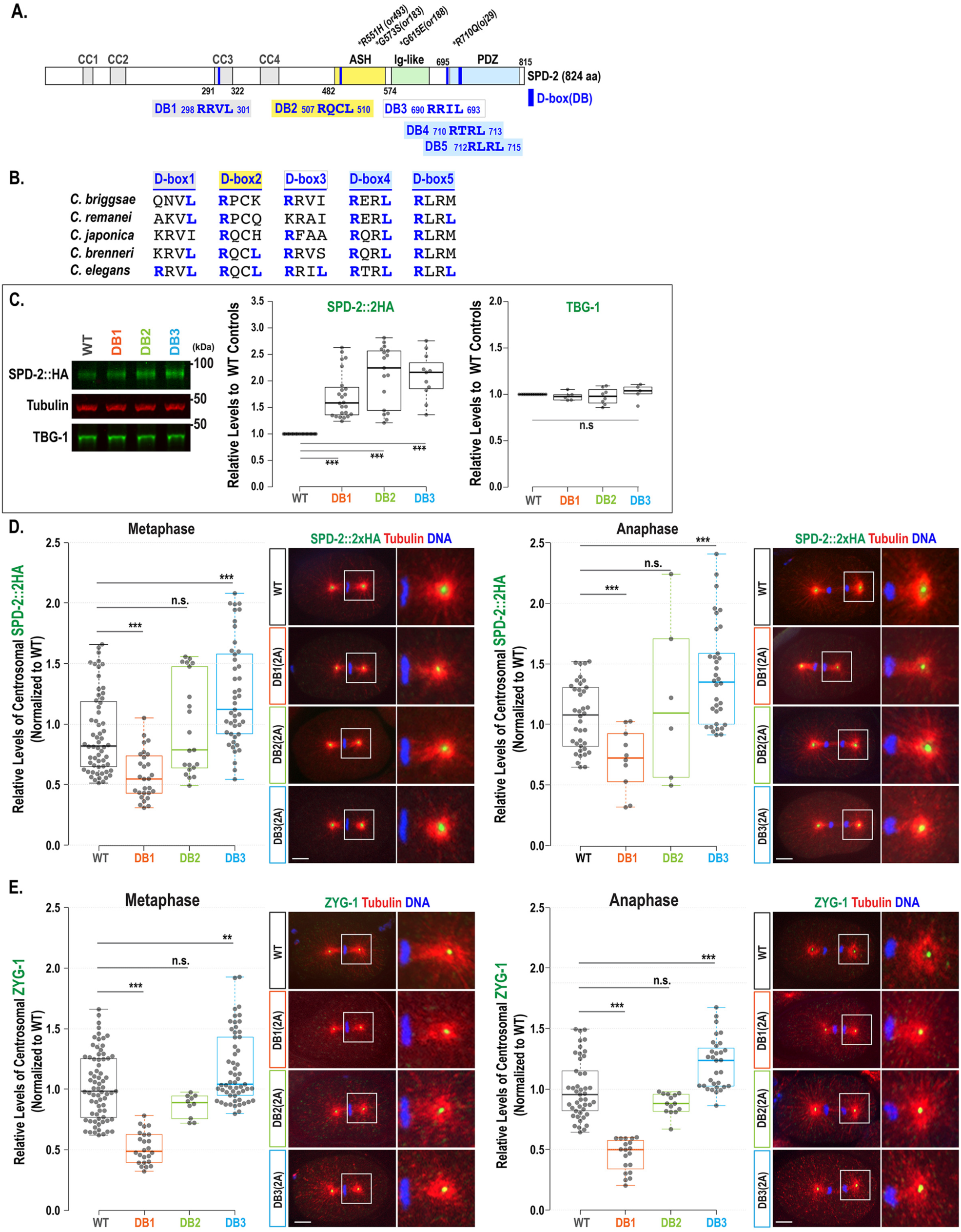
Mutating D-box motifs elevates overall cellular levels of SPD-2. (**A**) The SPD-2 protein structure illustrates the locations of D-boxes (DB:1 through 5), functional domains, and mutant alleles. (**B**) Sequence alignments of D-box motifs in the *Caenorhabditis* species. (**C**) (Left) Representative western blot with anti-HA and anti-TBG-1 using embryonic lysates from individual degron mutants and wild-type (WT) controls expressing SPD-2::2xHA. Tubulin was used as a loading control. (Middle) Quantitative western blot analysis shows significantly increased levels of SPD-2::2xHA in all three degron mutants. (Right) In contrast, there were no significant differences in TBG-1 levels among the degron mutants and wild-type controls. (**D-E**) (Left) Quantification of total centrosomal SPD-2::2xHA levels during the first mitotic cycle. Each dot represents a centrosome. (Right) Embryos stained for (**D**) SPD-2::2xHA and (**E**) ZYG-1 at the first mitotic metaphase and anaphase, with magnified images of posterior centrosomes. Bars, 10 μm. (**C-E**) In the plots, the box ranges from the first through the third quartile of the data. The thick bar indicates the median. The solid grey line extends 1.5 times the interquartile range or to the minimum and maximum data points. ^ns^*p*>0.05, \*\**p*<0.01, \*\*\**p*<0.001 (two-tailed t-tests).

We first examined whether APC/C^FZR-1^ targets SPD-2 by directly recognizing the canonical D-box degron. To determine if any of the five putative D-box motifs function as degrons *in vivo* by mediating FZR-1 and SPD-2 interaction, we used CRISPR/Cas9 genome editing to generate degron mutants by substituting two key residues within each D-box motif (RxxL) with alanine (AxxA) at their endogenous loci. Viable degron mutants were successfully produced for D-box1, D-box2, and D-box3, each with two-alanine substitutions, referred to as the SPD-2:DB1(2A), SPD-2:DB2(2A), and SPD-2:DB3(2A) mutations (**Table 1**). However, homozygous 2A mutations in D-box4 and D-box5 were lethal, indicating that the normal function of these motifs is essential for viability. Notably, a genome-wide genetic screen identified the *spd-2(oj29: R710Q)* mutation, which alters the first Arg (R) residue of D-box4 (710-RTRL-713) and causes centrosome duplication failure and embryonic lethality (**Fig. 2A**, Kemp et al. 2004). As a result, we were unable to evaluate the functional roles of the D-box4 and D-box5 degron motifs *in vivo*. Due to this limitation, we focused our analysis on D-boxes 1, 2, and 3 (DB1, 2, and 3) using viable degron mutants to assess their functional effects in this study.

**Table 1.**
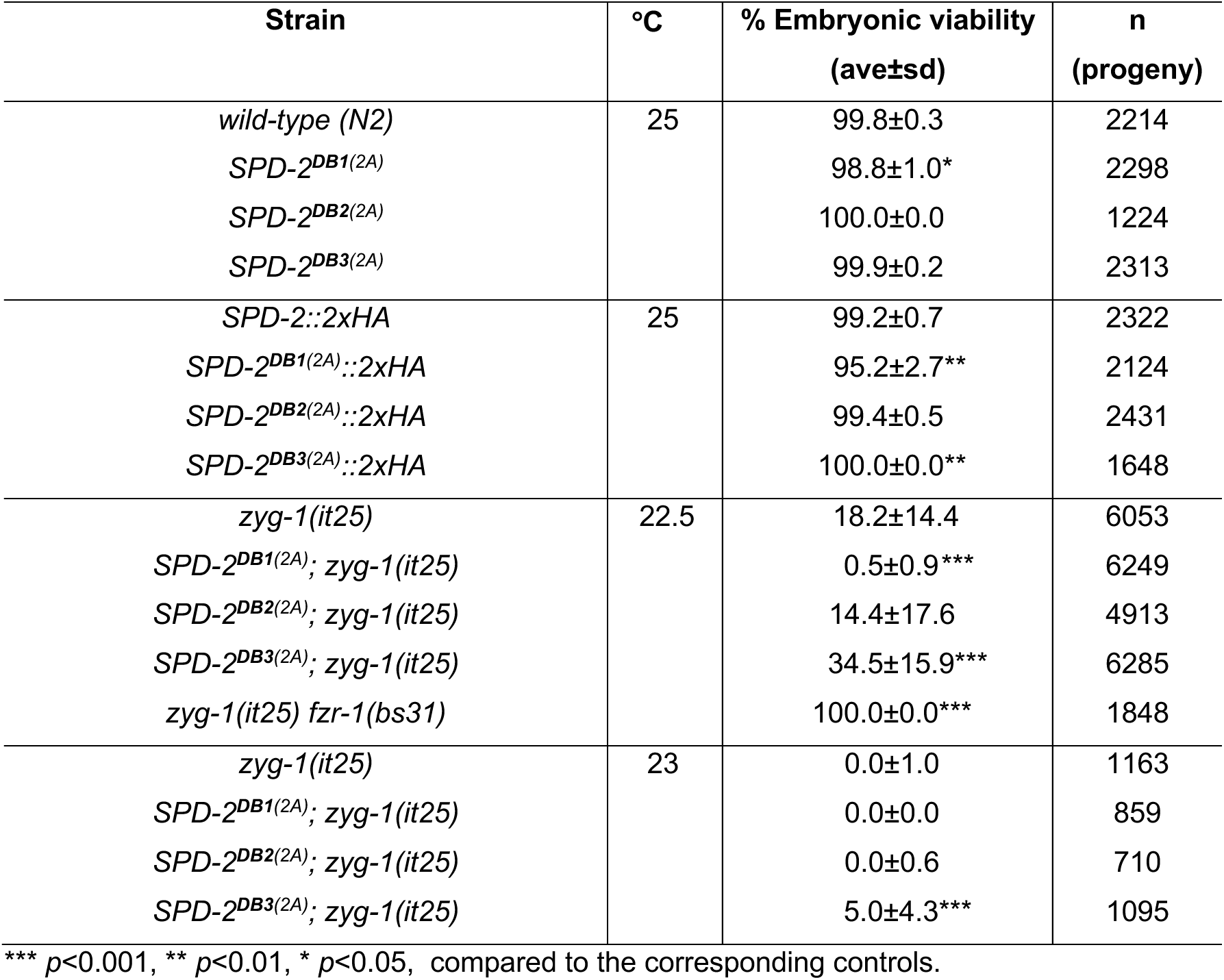
Genetic Analysis.

### Mutating the D-box leads to increased overall cellular SPD-2 levels

If APC/C^FZR-1^ recognizes SPD-2 through its D-box motif, then mutating the D-box degron should inhibit APC/C^FZR-1^ binding to SPD-2, preventing its degradation and resulting in higher SPD-2 protein levels. To assess how the D-box motif affects overall SPD-2 levels, we conducted quantitative Western blot analysis using protein lysates from each degron mutant embryo expressing SPD-2::2xHA (Medley et al. 2023) (**Fig. 2C**). Our results show that SPD-2 levels in all three D-box mutant embryos are significantly higher (DB1: 1.72±0.43-fold, n=24; DB2: 2.10±0.57-fold, n=17; DB3: 2.09±0.43-fold, n=11, *p*<0.001) compared to wild-type embryos (1.00±0.0-fold, n=17). In contrast, TBG-1 levels remain unaffected in degron mutants. These findings suggest that mutating the D-box motifs renders SPD-2 resistant to degradation, supporting the idea that the three D-box motifs (DB1, DB2, and DB3) in SPD-2 function as degrons directly recognized by APC/C^FZR-1^. Blocking this recognition prevents SPD-2 degradation, leading to its accumulation and mimicking SPD-2 overexpression.

### Only the SPD-2:DB3(2A) mutation results in elevated centrosomal SPD-2 levels

Quantitative Western blot shows that the SPD-2 degron mutations lead to increased total SPD-2 levels in the embryo, consistent with enhanced protein stabilization. To assess whether this stabilization influences SPD-2 levels at centrosomes, we examined how each degron mutation changes centrosomal SPD-2 levels. We performed quantitative immunofluorescence (IF) to compare centrosome-associated SPD-2 signals during the first mitosis in embryos expressing SPD-2::2xHA, stained with anti-HA tag and anti-tubulin (Medley et al. 2021; **Fig. 2D**).

The DB3(2A) mutation resulted in elevated centrosomal SPD-2 levels (1.35±0.43-fold, *p*<0.01, n=45 at metaphase; 1.32±0.42-fold, *p*<0.0001, n=34 at anaphase) compared to WT controls (1.00±0.33-fold, n=60 at metaphase; 1.00±0.27-fold, n=38 at anaphase), indicating that mutating the D-box3 does not impair centrosomal localization. Instead, the increased centrosomal signal likely reflects the elevated pool of cellular SPD-2 caused by protein stabilization. In contrast, the DB1(2A) mutation caused a significant decrease in centrosomal SPD-2 (0.59±0.20-fold, *p*<0.001, n=28 at metaphase; 0.66±0.26-fold, *p*<0.001, n=10 at anaphase) compared to controls. This dramatic reduction of centrosomal SPD-2 in the DB1(2A) mutant indicates that the D-box1 motif is critical for SPD-2 localization to centrosomes, implicating its dual role as both a degron and a key site for subcellular localization. Meanwhile, the DB2(2A) mutation caused no significant changes in centrosomal SPD-2 (1.06±0.40-fold, *p*=0.55, n=21 at metaphase; 1.12±0.68-fold, *p*=0.40, n=6 at anaphase) although the DB2(2A) mutation increases overall SPD-2 levels, indicating that the SPD-2:DB2(2A) mutation partially impairs SPD-2 localization to centrosomes. These degron mutants also showed consistent trends in centriolar SPD-2 levels (Supplementary **Fig. 2A**). The DB3(2A) mutant embryos exhibit higher levels (1.23±0.28-fold, n=31 at metaphase; 1.13±0.22-fold, n=35 at anaphase), while the DB1(2A) mutants have reduced levels (0.57±0.18-fold, n=28 at metaphase; 0.63±0.17-fold, n=10 at anaphase) and the DB2(2A) mutants show no significant change (1.07±0.46-fold, n=21 at metaphase; 0.98±0.2-fold, n=8 at anaphase), compared to WT controls (1.00±0.33-fold, n=56 at metaphase; 1.00±0.23-fold, n=27 at anaphase).

Furthermore, to assess the impact of SPD-2 degron mutations on ZYG-1 recruitment, we quantified centrosomal ZYG-1 levels in the degron mutants stained with anti-ZYG-1 and anti-tubulin (**Fig. 2E**). Consistent with their elevated SPD-2 levels, DB3(2A) mutants exhibited a significant increase in centrosomal ZYG-1 (1.14±0.29-fold, *p*<0.01, n=57 at metaphase; 1.21±0.21-fold, *p*<0.001, n=31 at anaphase) compared to WT controls (1.00±0.28-fold, n=72 at metaphase; 1.00±0.25-fold, n=45 at anaphase). In contrast, DB1(2A) mutants, which display reduced SPD-2 at centrosomes, showed a marked decrease in centrosomal ZYG-1 (0.50±0.13-fold, *p*<0.001, n=24 at metaphase; 0.45±0.13-fold, *p*<0.001, n=19 at anaphase). DB2(2A) mutants exhibited a slight reduction with no significant difference (0.84±0.10-fold, *p*=0.07, n=10 at metaphase; 0.88±0.09-fold, *p*=0.09, n=14 at anaphase). These results align with the observed patterns of centrosomal SPD-2 levels across the degron mutants (**Fig. 2D**). These data reveal a strong correlation between centrosomal SPD-2 levels and ZYG-1 recruitment: mutants with elevated centrosomal SPD-2 recruit more ZYG-1 to centrosomes, while those with reduced centrosomal SPD-2 show a proportional decrease in ZYG-1.

### The SPD-2:DB3(2A) mutation leads to *zyg-1* suppression

Next, we investigated how specific degron mutations affected SPD-2 activity in regulating centrosome assembly, using the hypomorphic *zyg-1(it25)* genetic background. The *zyg-1(it25)* allele carries a temperature-sensitive (ts) mutation (P442L) in the cryptic polo-box (CPB) domain, which is critical for centrosomal recruitment of ZYG-1 (Kemphues et al. 1988; Shimanovskaya et al. 2014). At the restrictive temperature 24°C, *zyg-1(it25*) mutant embryos fail to duplicate centrosomes during the first mitosis, leading to monopolar spindle formation in the second mitosis and resulting in 100% embryonic lethality (O’Connell et al. 2001). In the molecular hierarchy, SPD-2 acts upstream of ZYG-1 and is required for ZYG-1 loading to centrioles before procentriole formation (Pelletier et al. 2006).

If APC/C^FZR-1^ targets SPD-2 for degradation through its D-box degron, then mutating the D-box motif should block APC/C^FZR-1^ binding, thus stabilizing SPD-2. Consequently, hyper-stabilized SPD-2, similar to overexpression, could compensate for the reduced ZYG-1 activity in *zyg-1(it25)* mutants. To determine whether any SPD-2 D-box mutation could rescue *zyg-1(it25)* phenotypes, we introduced each mutation into the *zyg-1(it25)* background by genetic crossing and examined their effects on embryonic survival (**Fig. 3A**, **Table 1**) and centrosome duplication (**Fig. 3B**). Consistent with our hypothesis, the DB3(2A) mutation restored embryonic viability (35±16% vs 18±14%, *p*<0.001) and bipolar spindle formation (61±15% vs 34±22%, *p*<0.01) in *zyg-1(it25)* mutants, compared to *zyg-1(it25)* controls, at the semi-restrictive temperature of 22.5°C. This rescue effect was more potent at 22.5°C (34.5±15.9%) than at a more restrictive temperature of 23°C (5.0±4.3%; **Table 1**), suggesting that the SPD-2:DB3(2A) mutation depends on residual ZYG-1 activity for suppression. However, suppression of *zyg-1* by the SPD-2:DB3(2A) mutation appeared significantly weaker than that observed with the *fzr-1* mutation (100% bipolar spindles and embryonic viability; **Fig. 3B**, **Table 1**) (Medley et al. 2021).

**Figure 3.**
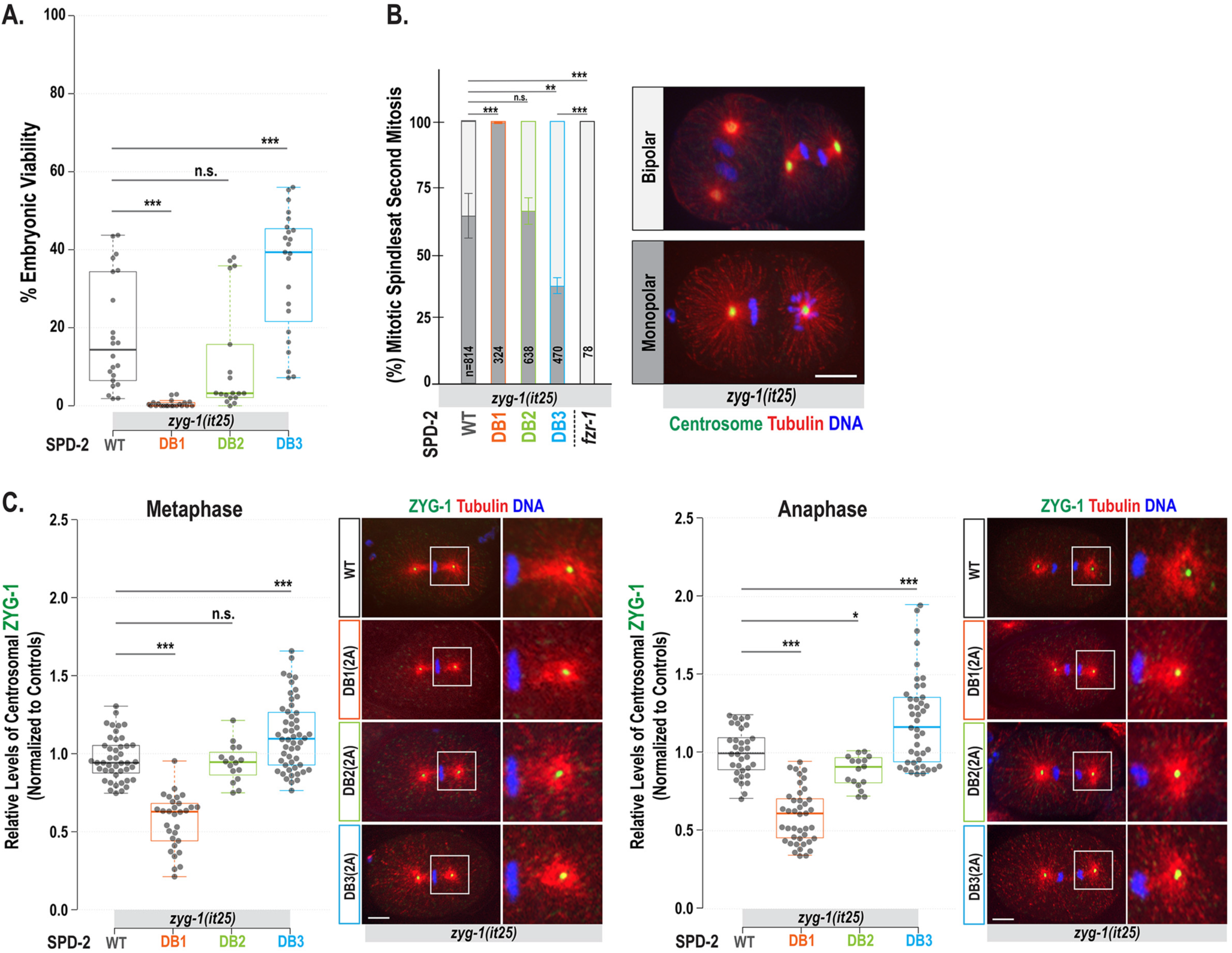
Mutating SPD-2 D-box3 leads to the suppression of *zyg-1* phenotypes and increased centrosomal ZYG-1 levels. (**A**) Embryonic viability at 22.5°C (See **Table 1**). Each dot represents a hermaphrodite. (**B**) Quantification of monopolar (dark grey) and bipolar (light grey) spindles in two-cell stage embryos at 23°C. Average and standard deviation (SD) are presented. n is the number of blastomeres. Embryos stained for centrosomes (TBG-1), microtubules, and DNA, illustrate mitotic spindles at the second mitosis. (**C**) (Left) Quantification of centrosome-associated ZYG-1 levels at the first mitotic metaphase and anaphase in *zyg-1(it25)* mutant backgrounds with individual degron mutations. Each dot represents a centrosome. (Right) Embryos stained for ZYG-1 and microtubules during the first mitotic cycle, with magnified images of posterior centrosomes. (**B-C**) Bars, 10 μm. (**A,C**) In the plots, the box ranges from the first through the third quartile of the data. The thick bar indicates the median. A solid grey line extends 1.5 times the interquartile range, or to the minimum and maximum data points. (**A-C**) ^ns^*p*>0.05, \**p*<0.05, \*\**p*<0.01, \*\*\**p*<0.001 (two-tailed t-tests).

In contrast, the DB1(2A) mutation markedly decreased embryonic viability (0.5±0.9% vs 18±14%, *p*<0.001; **Fig. 3A**, **Table 1**) and bipolar spindles (1.2±2.2% vs 34±22%, *p*<0.001; **Fig. 3B**) in *zyg-1(it25)* mutants, although the DB1(2A) mutant alone remained largely viable (99.2±1.1%; **Table 1**). These results indicate that *zyg-1(it25)* provides a sensitized genetic background where SPD-2:DB1(2A) causes near-complete lethality, demonstrating a strong synthetic effect. Similarly, although *spd-2* and *zyg-1* mutations are individually recessive, *spd-2/+; zyg-1/+* double heterozygotes exhibit ∼75% embryonic lethality (Song et al. 2011). DB1(2A) mutant embryos show centrosomal SPD-2 levels at 59% of those in wild-type embryos, comparable to those in *spd-2* heterozygotes (*spd-2*/+), despite exhibiting elevated cellular SPD-2 levels. Thus, D-box1 appears to regulate centrosomal loading of SPD-2 independently of its degron function. Although 59% of centrosomal SPD-2 is sufficient to support ZYG-1 loading and normal centrosome duplication in a wild-type background, combining the SPD-2:DB1(2A) mutation with homozygous *zyg-1(it25)* severely disrupts centrosome duplication and mitosis, leading to near-complete embryonic lethality at the semi-restrictive temperature (22.5 °C), where *zyg-1(it25)* alone causes ∼80% lethality (**Table 1**). These findings suggest that the D-box1 motif promotes proper localization of SPD-2 at centrosomes and may facilitate its interaction with ZYG-1. D-box1 resides within the coiled-coil (CC3) domain (**Fig. 2A**), and the DB1(2A) mutation consists of two alanine substitutions (R298A and L301A). Since coiled-coil domains are known to mediate protein–protein interactions (Murph et al. 2022), these substitutions likely impair the ability of SPD-2 to interact with binding partners necessary for centrosome function.

Meanwhile, the DB2(2A) mutation had a minimal negative impact on embryonic viability (14.4±17.6% vs 18±14%, *p*=0.36; **Table 1**, **Fig. 3A**) and bipolar spindle formation (30.2±18.5% vs 34±22%, *p*=0.55; **Fig. 3B**) in *zyg-1(it25)* mutants. Like DB1(2A), the DB2(2A) mutation increased overall SPD-2 levels but did not enhance its centrosomal activity, suggesting a partial defect in SPD-2 loading, though much milder than DB2(2A). Since it resides within the ASH domain, which is known to target SPD-2 to centrosomes (Murph et al. 2022), this may partly explain the modest yet negative impact of the DB2(2A) mutation on embryonic viability, unlike its effect on SPD-2 stability.

### Increased centrosomal ZYG-1 levels in SPD-2:DB3(2A) mutants

In *zyg-1(it25)* embryos, centrosomal ZYG-1 is reduced to ∼40% of WT controls (Medley et al. 2017), impairing the recruitment of SAS-5 and SAS-6, which are required for subsequent SAS-4 loading to centrosomes (Pelletier et al. 2006). Since SPD-2 acts upstream of ZYG-1 and promotes centrosomal recruitment of ZYG-1, the observed suppression of *zyg-1* mutant phenotypes likely results from increased centrosomal ZYG-1 levels caused by SPD-2 stabilization. Consequently, increased levels of centrosomal ZYG-1 may restore the recruitment of downstream factors, rendering centrosomes competent to assemble new centrioles in *zyg-1(it25)* mutants.

To assess how the SPD-2 degron mutation influenced ZYG-1 recruitment in *zyg-1(it25)* mutants, we measured centrosomal ZYG-1 levels in *zyg-1(it25)* embryos carrying each degron mutation (**Fig. 3C**). As expected, the SPD-2:DB3(2A) mutation led to increased centrosomal ZYG-1 levels (1.15±0.22-fold, *p*<0.001, n=52 at metaphase; 1.20±0.29-fold, *p*<0.001, n=43 at anaphase) compared to *zyg-1(it25)* controls (1.00±0.14-fold, n=44 at metaphase; 1.00±0.15-fold, n=33 at anaphase). In contrast, the DB1(2A) mutation significantly decreased centrosomal ZYG-1 (0.53±0.14-fold, *p*<0.001, n=24 at metaphase; 0.31±0.06-fold, *p*<0.001, n=6 at anaphase), while DB2(2A) mutants exhibited a slight decrease (0.92±0.18-fold, *p*=0.466, n=12 at metaphase; 0.91±0.29-fold, *p*=0.02, n=10 at anaphase). These results align with the patterns of centrosomal SPD-2 levels across each degron mutant combined with *zyg-1(it25)* (Supplementary **Fig. 3A**). The SPD-2:DB3(2A) mutation caused higher centrosomal SPD-2 levels (1.22±0.35-fold, *p*<0.001, n=66 at metaphase; 1.25±0.28-fold, *p*<0.001, n=69 at anaphase), while DB1(2A) resulted in a dramatic decrease (0.74±0.25-fold, *p*<0.001, n=22 at metaphase; 0.74±0.30-fold, *p*<0.001, n=33 at anaphase), and DB2(2A) showed a slight reduction (0.86±0.22-fold, *p*<0.01, n=42 at metaphase; 1.03±0.24-fold, *p*=0.67, n=13 at anaphase), relative to *zyg-1(it25)* controls (1.00±0.29-fold, n=76 at metaphase; 1.00±0.19-fold, n=30 at anaphase). These findings indicate that the SPD-2:DB3(2A) mutation leads to increased centrosomal ZYG-1, which enhances the recruitment of downstream factors and restores centrosome duplication and embryonic viability in *zyg-1* mutants. This suggests a possible mechanism by which SPD-2:DB3(2A) suppresses the *zyg-1* phenotype.

Finally, we observed that the SPD-2:DB3(2A) mutation results in lower centrosomal SPD-2 levels than the *fzr-1(bs31)* mutation in *zyg-1(it25)* mutant backgrounds (1.25±0.28-fold, n=69 vs 2.59±0.89-fold, n=14, *p*<0.001; Supplementary **Fig. 3B**), consistent with the weaker suppression of the *zyg-1* phenotype by the SPD-2:DB3(2A) mutation relative to the *fzr-1* mutation. SPD-2:DB3(2A) mutant embryos show increased cellular SPD-2 levels, resulting in higher centrosomal levels of SPD-2 and ZYG-1. However, the SPD-2:DB3(2A) mutation alone produces a much weaker rescue of embryonic viability (1.25±0.28% vs 100%, **Table 1**) and bipolar spindle formation (60±15% vs 100±0%, **Fig. 2B**) in *zyg-1(it25)* mutants compared to the *fzr-1* mutation. Similar results were observed in our earlier studies, where ZYG-1 or SAS-5 degron mutations led to *zyg-1* suppression that was less effective than the *fzr-1* mutation (Medley et al. 2017; Medley et al. 2021). Several factors may account for the relatively weak suppression of *zyg-1* by the DB3(2A) single mutation. First, we were unable to test the *in vivo* functions of D-box4 and D-box5 because their 2A substitutions are homozygous lethal, suggesting that these degrons may play a more dominant role in SPD-2 turnover than D-box3. Second, we could not assess the combined effects of D-boxes 1 through 3 on *zyg-1* suppression because D-box1 and D-box2 appear to be required for centrosomal localization of SPD-2 and fail to rescue *zyg-1* phenotypes, making triple mutant analysis challenging. Thus, the DB3(2A) mutation alone is unlikely to entirely block APC/C^FZR-1^-dependent degradation of SPD-2, resulting in only a partial rescue of the *zyg-1(it25)* phenotype, which is weaker than the *fzr-1* mutation. Another possibility is that APC/C^FZR-1^ targets additional centrosome proteins upstream of SPD-2, such as SAS-7 (Sugioka et al. 2017). In this scenario, loss of APC/C^FZR-1^ would enhance ZYG-1 recruitment both by stabilizing ZYG-1 directly and by increasing the levels of SPD-2 and other upstream factors, such as SAS-7, required for ZYG-1 loading. However, testing the contribution of SAS-7 is beyond the scope of the present study and will require additional investigation.

### SPD-2:DB1(2A) mutant embryos exhibit abnormal cell division patterns

During our genetic analysis at 25°C, we observed a low but significant rate of embryonic lethality in the SPD-2:DB1(2A) mutants, and this lethality was further exacerbated in the SPD-2:DB1(2A) mutant embryos tagged with 2xHA (1.2% vs 4.8% lethality, **Table 1**). To investigate the cause of embryonic lethality, we examined SPD-2:DB1(2A):2xHA mutant embryos stained for centrosomes, microtubules, and DNA (**Fig. 4**). At 25°C, DB1(2A) mutant embryos display abnormal cell division phenotypes. Consistent with reduced levels of SPD-2 (**Fig. 2D**) and ZYG-1 (**Fig. 2E**) at centrosomes, these embryos display tiny and often highly asymmetric centrosomes during the first mitotic metaphase. Such small and asymmetric centrosomes have also been observed in ZYG-1:4D mutant embryos, which exhibit unstable and reduced levels of centrosomal ZYG-1, indicating a partial block in centrosome duplication (Medley et al. 2023). DB1(2A) mutants also display abnormal patterns in DNA segregation, cell division timing, and mitotic division patterns, compared to the WT control, although at a low frequency (<5%, n>100). These cell division defects likely account for embryonic lethality and further confirm that the D-box1 motif functions not only as a degron motif but also plays a critical role in SPD-2 localization and its centrosomal activity. Therefore, our study identified new key sites essential for SPD-2 activity in regulating centrosome assembly during early embryonic development.

**Figure 4.**
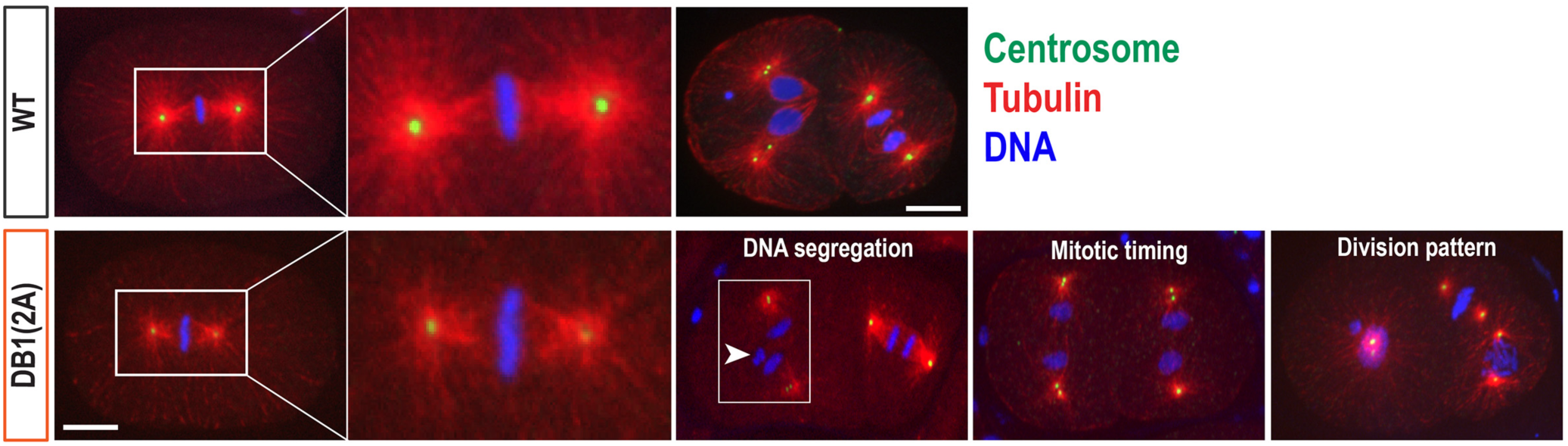
SPD-2:DB1(2A) mutants exhibit cell division defects during early embryogenesis. Embryos expressing SPD-2::2xHA stained for centrosomes (SAS-4), microtubules, and DNA. At 25°C, SPD-2:DB1(2A) mutant embryos exhibit abnormal cell division phenotypes, including small centrosomes and highly asymmetric centrosome sizes during the first mitotic metaphase, DNA missegregation (arrowhead), synchronous mitotic timing, and aberrant patterns of mitotic divisions, compared to the WT embryos (Top). Bar, 10 μm.

In summary, this study reports that *C. elegans* SPD-2 is a direct substrate of APC/C^FZR-1^ and illustrates a regulatory network that links protein degradation, centrosomal localization, and interactions with other partners. We showed that SPD-2 physically interacts with FZR-1 *in vivo*, and that loss of APC/C^FZR-1^ results in elevated levels of both cellular and centrosomal SPD-2. These results expand the repertoire of APC/C^FZR-1^ substrates beyond SAS-5 and ZYG-1 (Medley et al. 2017; Medley et al. 2021), highlighting the crucial role of APC/C^FZR-1^ in controlling centrosome assembly during *C. elegans* embryogenesis. Importantly, our findings also suggest that APC/C^FZR-1^ may target additional centrosome factors, with SAS-7 (Sugioka et al. 2017) as a strong candidate for future investigation.

Our degron motif analysis reveals both canonical and noncanonical functions. While all three D-box mutations (DB1, DB2, and DB3) stabilize SPD-2, only the DB3 mutation restores centrosome duplication and embryonic viability in *zyg-1* mutants. Our results also reveal that D-box1 and D-box2 have functions beyond their roles as degron motifs. The D-box1 is required for efficient centrosomal localization of SPD-2 and may facilitate its interaction with ZYG-1. Consistently, DB1(2A) mutants show reduced centrosomal SPD-2 and ZYG-1 levels, leading to monopolar spindle formation and increased embryonic lethality in *zyg-1* mutants. This suggests that D-box1 is important for both stabilizing and centrosomal targeting of SPD-2. By contrast, DB2 mutation increased total SPD-2 levels but did not alter centrosomal pools or rescue *zyg-1* phenotypes. These findings demonstrate that degron motifs serve multiple roles, with D-box1 and D-box2 involved in both proteasomal degradation and centrosomal functions.

Finally, our findings have broader evolutionary implications. SPD-2/CEP192 is a highly conserved key regulator of centrosome assembly in metazoans (Kemp et al. 2004; Pelletier et al. 2004; Zhu et al. 2008). Because dysregulation of centrosome factors is closely associated with genomic instability and tumorigenesis in humans (Nigg and Holland 2018), understanding the degron-dependent regulation of SPD-2/DSpd-2 in *C. elegans* and *Drosophila* (Meghini et al. 2016; Meghini et al. 2023) provides an evolutionary framework for investigating analogous mechanisms in human cells. Future work should explore how APC/C^FZR-1^ influences centrosome assembly and microtubule dynamics, with potential implications for human health and cancer biology.

## Data and Reagent Availability

The authors confirm that all data necessary to support the conclusions of the article are presented within the article, figures, tables, and supplementary material. Strains used in this study are available upon request.

## Acknowledgements

We thank former and current members of the Song lab for their technical support. We especially thank WormBase and the *Caenorhabditis* Genetics Center (CGC). WormBase is supported by grant U24HG002223 from the National Human Genome Research Institute at the US National Institutes of Health, the UK Medical Research Council, and the UK Biotechnology and Biological Sciences Research Council. The CGC (St. Paul, MN), is funded by the National Institutes of Health Office of Research Infrastructure Programs (P40 OD010440).

## Author Contributions

M.H.S. designed the experiments, wrote the manuscript, and performed quantifications of confocal imaging and protein levels from western blots. R.N.Y., J.D., P.R., and M.H.S. performed experiments and provided data.

