## Supplemental Files for "Distinct D-box Motifs in SPD-2 Mediate APC/C^FZR-1^-Dependent Degradation and Centrosomal Localization in *Caenorhabditis elegans* Embryos"

**Supplementary Table 1. List of *C. elegans* Strains**

| Strain | Genotype | Origin |
|---|---|---|
| **N2** | *wild-type* | CGC |
| **OC14** | *zyg-1(it25[ZYG-1$^{P442L}$]) II* | O'Connell et al., 2001 |
| **OC201** | *zyg-1(it25) fzr-1(bs31) II* | Kemp et al., 2007 |
| **MTU6** | *fzr-1(bs31) II* | Medley et al., 2017 |
| **MTU7** | *mat-3(or180) III* | Golden et al., 2000 |
| **MTU490** | *spd-2(mhs650[SPD-2$^{DB1(2A)}$]) I; dpy-10(cn64)/+ II* | This study |
| **MTU686** | *spd-2(mhs720[SPD-2$^{DB2(2A)}$]) I; dpy-10(cn64)/+ II* | This study |
| **MTU493** | *spd-2(mhs653[SPD-2$^{DB3(2A)}$]) I; dpy-10(cn64)/+ II* | This study |
| **MTU696** | *spd-2(mhs651[SPD-2$^{DB1(2A)}$]) I; zyg-1(it25) II* | This study |
| **MTU689** | *spd-2(mhs720[SPD-2$^{DB2(2A)}$]) I; zyg-1(it25) II* | This study |
| **MTU546** | *spd-2(mhs653[SPD-2$^{DB3(2A)}$]) I; zyg-1(it25) II* | This study |
| **MTU337** | *spd-2(mhs580[SPD-2::2xHA]) I; dpy-10(cn64)/+ II* | Medley et al., 2021 |
| **MTU369** | *fzr-1(mhs470[Ollas::FZR-1]) II* | Medley et al., 2021 |
| **MTU375** | *spd-2(mhs580[SPD-2::2xHA]) I; zyg-1(it25) II* | This study |
| **MTU806** | *spd-2(mhs580[SPD-2::2xHA]) I; fzr-1(mhs470[Ollas::FZR-1]) II* | This study |
| **MTU816** | *spd-2(mhs580[SPD-2::2xHA]) I; fzr-1(bs31) II* | This study |
| **MTU822** | *spd-2(mhs650mhs770[SPD-2$^{DB1(2A)}$::2xHA]) I* | This study |
| **MTU826** | *spd-2(mhs720mhs774[SPD-2$^{DB2(2A)}$::2xHA]) I* | This study |
| **MTU829** | *spd-2(mhs653mhs777[SPD-2$^{DB3(2A)}$::2xHA]) I* | This study |

**Supplementary Table 2. List of crRNA for CRISPR/Cas9 Genome**

| Gene | Target | Sequence (5'-3') |
|---|---|---|
| ***dpy-10*** (Arribere et al., 2014) | *dpy-10(cn64)* co-CRISPR | UUCUGCUGUCUUGAUUGACG |
| ***spd-2*** | C-terminus | UCUAUUCGAAAAUCUUGUAU |
| ***spd-2*** | D-box1 | AUGGUUCGAAGAGUGCUCAA |
| ***spd-2*** | D-box2 | UCGAACAAGACACUGACGAU |
| ***spd-2*** | D-box3 | CUCCAGGAAGAAUACGUCGC |
| ***spd-2*** | D-box4, 5 | UCAAAUUAACAAUCGAAGUC |

**Supplementary Table 3. List of ssODN Homologous Repair Templates for CRISPR/Cas9 Genome Editing**

| Gene | Variation | Sequence (5'-3') |
|---|---|---|
| ***dpy-10***<br>(Arribere et al., 2014) | *dpy-10(cn64)* | CACTTGAACTTCAATACGGCAAGATGAGAATGACT<br>GGAAACCGTACCGCATGCGGTGCCTATGGTAGCG<br>GAGCTTCACATGGCTTCAGACCAACAGCCTAT |
| ***spd-2*** | C-terminal 2xHA Tag | AATCAGACATTTGTCAACGACGTTACAATTGTTCC<br>GAATACAAGATTTTCGAATAGAAAGGGAGGTTCC<br>GGTGGATCTGGTGGATCCTACCCATACGATGTTC<br>CAGATTACGCTTATCCATATGATGTTCCAGATTAT<br>GCTTAAAATCTTAACCTAACTTTCCAAATATTCTCT<br>G |
| ***spd-2*** | D-box1(AxxA) | TGATGACAAATCAAACTATTAATGAGTCAATGGTT<br>GCTCGCGTCGCTAAAGGAAACAATAAGAATCAAG<br>ATCTGTTCGCAGCATTAG |
| ***spd-2*** | D-box2(AxxA) | TTGAGAGTTGAAGTCGAAGTGGAAAACATTTCCGA<br>TGCTCAGTGTGCTGTTCGAGCTAGTACGGATAGT<br>ACCACACCAGTTTATCAAATCCT |
| ***spd-2*** | D-box3(AxxA) | TCTTCGCTTTCGACAGCCTCAACAACATCCTCATT<br>CGAGGCTCGTATTGCTCCTGGAGCCAAATTCTTT<br>GTGCATGTCGTTTGGGG |
| ***spd-2*** | D-box4(AxxA) | TCTTTGTGCATGTCGTTTGGGGTGAAGAAACCATG<br>GCTACTAGAGCTCGATTGTTAATTTGAATACAGAT<br>GAAATGGTATT |
| ***spd-2*** | D-box5(AxxA) | GTGCATGTCGTTTGGGGTGAAGAAACCATGCGAA<br>CTGCTCTTCGAGCGTTAATTTGAATACAGATGAAA<br>TGGTATTACATTT |

### Supplementary Figures

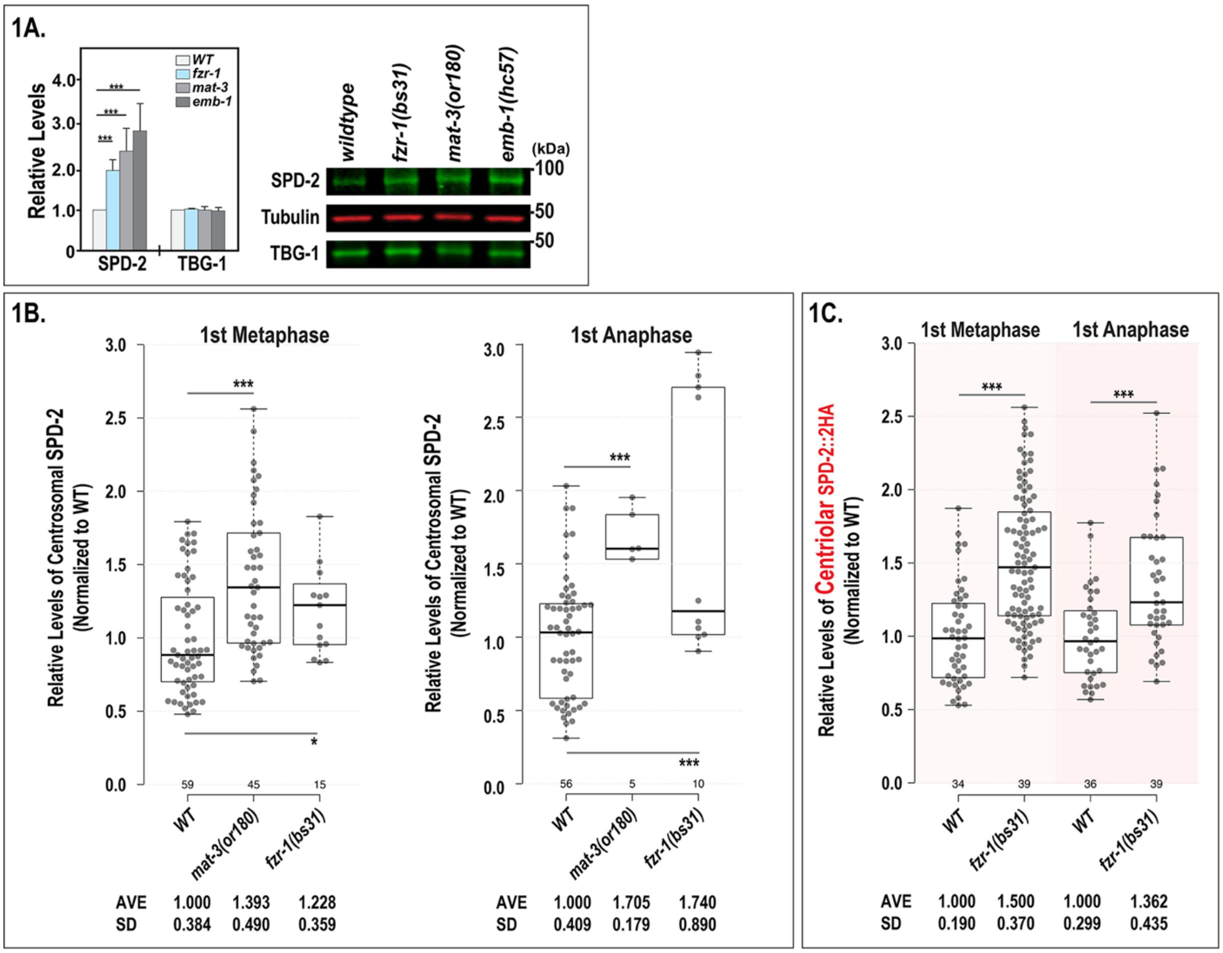


**Supplementary Figure 1.** (**A**) Overall SPD-2 levels are more than 2-folds higher in *fzr-1(bs31), mat-3(or180),* and *emb-1(hc57)* mutants. In contrast, TBG-1 levels remain unchanged across the strains. Representative Western blot with anti-SPD-2 and anti-TBG-1. Tubulin was used as a loading control. Average values are presented, and error bars are SD (n>5). ****p*<0.001 (two-tailed t-tests). (**B**) Like *fzr-1(bs31)* mutants, *mat-3(or188)* mutant embryos exhibit elevated levels of total centrosomal SPD-2::2xHA during the first mitosis compared to wildtype (WT) controls. (**C**) Quantification of centriolar SPD-2::2xHA levels in *fzr-1(bs31)* mutant embryos and WT controls at the first metaphase and anaphase. (**B-C**) Each dot represents a centrosome. In the plots, the box ranges from the first through the third quartile of the data. The thick bar indicates the median. Solid grey line extends 1.5 times the interquartile range or to the minimum and maximum data points. **p*<0.05*,* ***p*<0.01*,* ****p*<0.001 (two-tailed t-tests).

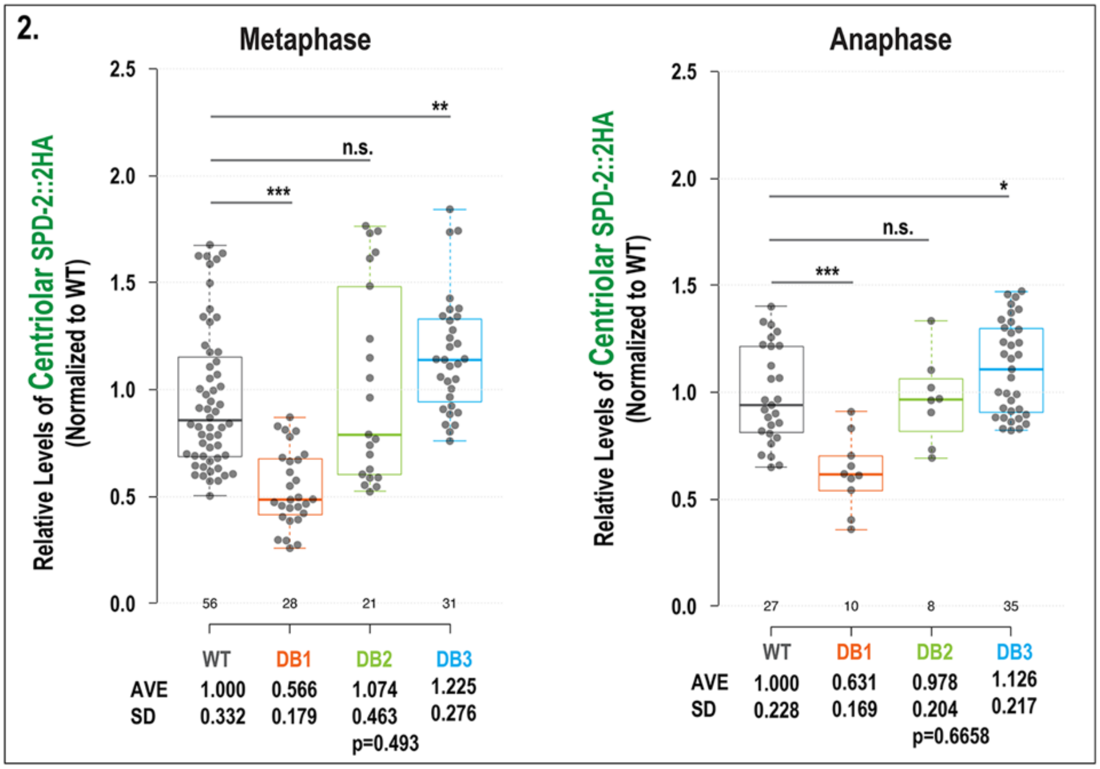


**Supplementary Figure 2:** Quantification of centriolar SPD-2::2xHA levels in SPD-2 degron mutants and WT controls at the first metaphase and anaphase. Each dot represents a centrosome. Average values are presented, and error bars are SD. In the plots, the box ranges from the first through the third quartile of the data. The thick bar indicates the median. Solid grey line extends 1.5 times the interquartile range or to the minimum and maximum data points. $^{ns}p>0.05$, $^{*}p<0.05$, $^{**}p<0.01$, $^{***}p<0.001$ (two-tailed t-tests).

Supplementary Data: Yim et al., 2025

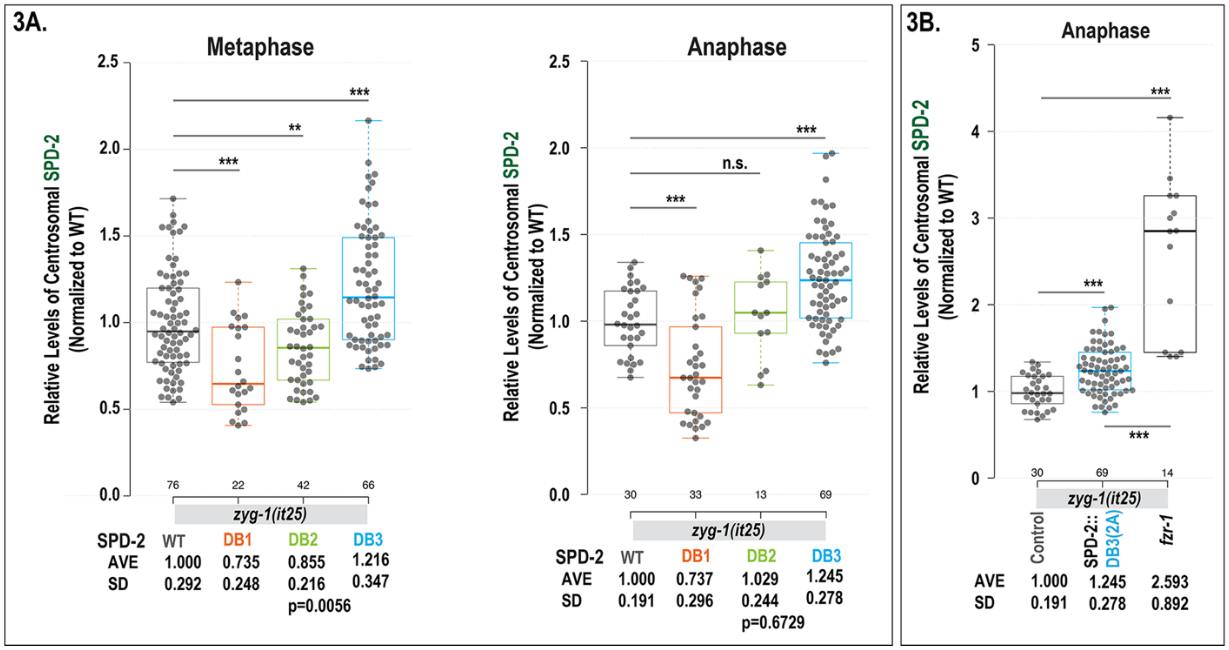


**Supplementary Figure 3.** (**A**) Quantification of total centromal SPD-2::2xHA levels during the first mitosis in *zyg-1(it25)* embryos carrying individual SPD-2 degron mutations or WT controls. (**B**) Quantification of total centrosomal SPD-2::2xHA levels in *zyg-1(it25)II* controls, *SPD-2::DB3(2A)I; zyg-1(it25)II* mutants, and *zyg-1(it25) fzr-1(bs31)II* embryos at the first anaphase. Each dot represents a centrosome. In the plots, the box ranges from the first through the third quartile of the data. The thick bar indicates the median. Solid grey line extends 1.5 times the interquartile range or to the minimum and maximum data point. $^{ns}p>0.05$, $^{**}p<0.01$, $^{***}p<0.001$ (two-tailed t-tests).
